# CAMOIP: A Web Server for Comprehensive Analysis on Multi-Omics of Immunotherapy in Pan-cancer

**DOI:** 10.1101/2021.09.10.459722

**Authors:** Anqi Lin, Ting Wei, Junyi Liang, Chang Qi, Mengyao Li, Peng Luo, Jian Zhang

**Author notes:** Correspondence may also be addressed to Peng Luo. Joint Authors. These authors have contributed equally to this work and share first authorship.

## Abstract

Immune checkpoint inhibitors (ICIs) have completely changed the therapeutic approach for tumor patients. Immunotherapy has also produced much needed data about mutation, expression, and prognosis, providing an unprecedented opportunity for discovering candidate drug targets and screening for immunotherapy-relevant biomarkers. Although existing web tools enable biologists to analyze the expression, mutation, and prognosis data on tumors, they are currently not able to carry out data mining and mechanism analyses related to immune checkpoint therapy. Thus, we developed our own web-based tool called Comprehensive Analysis on Multi-Omics of Immunotherapy in Pan-cancer (CAMOIP), in which we can screen prognostic markers and analyze the mechanisms involved with markers and immunotherapy (more than 4000 patients). The analyses include survival analysis, expression analysis, drug sensitivity analysis, mutational landscape, immune checkpoint analysis, immune related signature analysis, immune cell analysis, immune gene analysis, immunogenicity analysis and gene sets enrichment analysis (GSEA). This comprehensive analysis of biomarkers for immunotherapy can be carried out by a click of CAMOIP, and the software should greatly encourage the further development of immunotherapy. CAMOIP fills the gap between the big data of cancer genomics based on immunotherapy and providing comprehensive information to users, helping to release the value of current ICI-treated data resources. CAMOIP can be found in https://www.camoip.net.

## INTRODUCTION

Immunotherapy has dramatically changed the way clinicians approach the treatment of cancer. The introduction of immune checkpoint inhibitors (ICIs) has led to significant improvement in increasing immune activity and killing tumor cells(1). However, there is an increasing amount of clinical evidence that shows that only a small proportion of patients receive good clinical benefit in response to ICIs(2– 4). The exploration of biomarkers that can effectively distinguish immune responders from non-responders and screen potential beneficiaries is important for predicting the effect of immunotherapy and monitoring adverse immune related events(2).

In recent years, research on high-throughput sequencing of patients receiving ICIs treatment has produced considerable RNA-sequencing and mutation data, which provides an unprecedented opportunity for finding candidate drug targets and screening biomarkers relevant to immunotherapy(5–13). Examples of this data include the following: Samstein and his colleagues collected mutation data (via targeted sequencing) and clinical outcome data from pan-cancer patients treated with ICIs(5); Prat et al. collected the prognostic and expression data of pan-cancer patients after receiving PD-1 blockade(6); Rizvi et al. published a study on the association of high tumor mutation burden (TMB) with better clinical outcomes in NSCLC patients after receiving ICIs(7); Hwang and his colleagues published a study on NSCLC with therapeutic regimens of ICIs(8). In addition to pan-cancer and lung cancer data, the number of retrospective studies on the response to ICIs by melanomas is also considerable. For example, Allen et al. published clinical prognosis data and mutation data via whole exome sequencing (WES) on melanomas treated with ICIs (Cytotoxic T lymphocyte associate protein-4blockade)(9). In addition, Ulloa-Montoya et al.(10), Hugo et al.(11), Auslander et al.(12) and Riaz et al.(13) have published relevant research on melanoma patients receiving ICIs treatment and provided substantial clinical prognosis, mutation and/or expression data.

Although the rapid development of precision medicine such as immunotherapy based on ICIs has changed the therapeutic approach to tumor patients, the identification of suitable ICI-relevant biomarkers remains a challenge(5, 6, 14). Ideally, researchers would be able to utilize biomarkers to predict the efficacy of ICIs through genome-wide sequencing and multi-omics analysis, building a biomarker-based prediction model to comprehensively evaluate the tumor immune status of patients and formulate an individualized and precise combined treatment strategy(5, 15–18). Several studies support the potential value of biomarkers of ICIs, including those that have shown that: 1) the higher the expression level of programmed cell death-ligand 1 (PD-L1) in tumor cells, known as the tumor proportion score (TPS)(15), the higher the response rate to PD-L1 inhibitors(16); 2) the higher the CD8+ tumor infiltrating lymphocytes (TILs) score, the higher the clinical benefit in response to immunotherapy(17); 3) the expression level of the T cell infiltration gene expression profile was positively correlated with the clinical benefit of tumor immunotherapy(19); 4) the level of TMB was strongly correlated with the objective response rate (ORR) of PD-1/PD-L1 inhibitors(5, 18), and TMB level was positively correlated with the overall survival (OS) of immunotherapy patients(5). Despite the breadth of research, the above markers have their limitations. For example, the lack of standardization and consistency in PD-L1 detection methods, the heterogeneity of TMB in different laboratories and platforms, and the evaluation of next generation sequencing (NGS) being based on different panels(20–24). This suggests that the search for new biomarkers for predicting the efficacy of immunotherapy is an urgent problem still to be solved in the field of immunotherapy.

Currently, Xena(25), cBioPortal (http://www.cbioportal.org/)(26), Human Protein Atlas portal (HPA)(27), Expression Atlas(28), Gene Expression Profiling Interactive Analysis (GEPIA)(29), and GEPIA2(30) provide many useful visualizations and tools for gene expression analysis. Although these web tools are extremely valuable and widely used, they are unable to screen prognostic markers and provide multi omic mechanism analysis for immunotherapy cohorts. None of the existing tools can screen for predictive markers, nor provide survival analysis, GSEA, gene mutation map, expression analysis, drug sensitivty analysis, immune checkpoint analysis, immune related signature analysis, immune cell analysis, immune function gene analysis, and immunogenicity analysis for these cohorts. Based on the above unresolved requirements, we developed Comprehensive Analysis on Multi-Omics of Immunology in Pan-cancer (CAMOIP), a web-based tool that provides fast and customizable solutions to complement the existing tools. CAMOIP can be found in https://www.camoip.net.

## MATERIAL AND METHODS

### Implementation

The CAMOIP website is free for all users, and no login is required to access all functions. It is built with shiny R package(31), including ui.R, server.R and function.R. Server side and interactive data processing is performed by R script. The data table of the CAMOIP web site is displayed by DT:: datatableoutput, which allows users to query, select, and download data. Images are then generated by renderplot (Figure 1A) and can be downloaded in portable document format (PDF) and portable network graphics (PNG) images. All the pictures in CAMOIP are based on R software (v.4.0).

**Figure 1.**
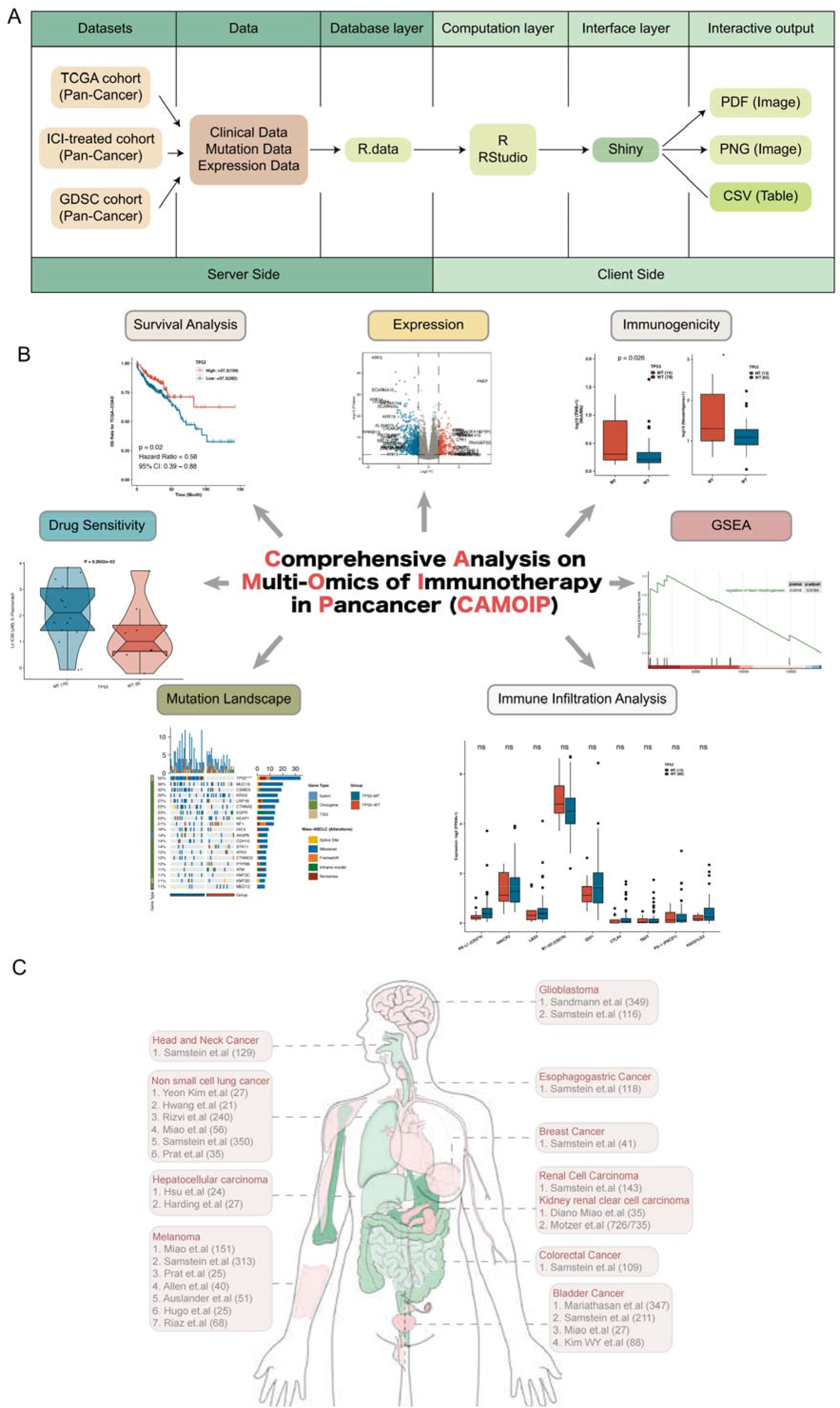
A. Schema describing data processing and data display for the CAMOIP visualization tool. B. The Home tab provides users with the details of datasets including ICI-treated and TCGA cohorts. C. The overview of the datasets in this study.

In order to establish a clinical cohort representing patients receiving ICIs, we collected data from cBioPortal(26) and Gene Expression Omnibus (GEO)(32). Since only one cohort with both mutation data and expression data was available, we used TCGAbiolinks R package(33) to download RNA-seq (raw count) with reference genome hg38 and mutation data measured by WES from the Genomic Data Commons (GDC) data portal (https://gdc.cancer.gov/about-data/publications/pancanatlas)(34) to further explore the relationship between selected biomarkers and cancer progression. After standardizing the raw count, fragments per kilobase of exon model per million mapped fragments (FPKM) was obtained. In addition, mutation and drug sensitivity data of tumor cell lines were obtained from the Genomics of Drug Sensitivity in Cancer (GDSC) database(35), with all data being stored on the server in the form of R.data. All R packages were detailed in the Table S1.

The function of the CAMOIP web server is divided into seven main tabs (Figure 1B): KM plotter, Expression, Drug Sensitivity, Mutational Landscape, Immune Infiltration, and immunity and GSEA. It provides key interactive functions corresponding to survival analysis, drug sensitivity analysis of tumor cells, an overview of gene mutation in patients, tumor immune microenvironment analysis, tumor immunogenicity analysis and customizable GSEA mapping.

## RESULTS

### Home

CAMOIP provides a graphical interface that briefly introduces all clinical cohort information used in this project (Figure 1C; Table S2). Users are able to find the sample number of the immunotherapy cohort corresponding to each tumor in the body diagram provided by the interface.

### Kaplan-Meier plotter

CAMOIP can be used for survival analysis according to either gene mutation level or gene expression level (Figure 2).This function allows users to select a tumor type and perform survival analysis for specific genes (grouped based on mutation or expression level) in either the immunotherapy or the The Cancer Genome Atlas (TCGA) cohort. The mutation levels can be further divided into wildtype (WT) and mutant (MT) based on all mutation; into WT and MT under non-synonymous mutation; into alternation (Alter) and non-alternation (Non-Alter) under alternation status; into best critical point group and the custom group proportion group at the expression level. For example, for OS based survival analysis of the Miao cohort, users can select non-small cell lung cancer (NSCLC) to group the WT and MT of the EGFR gene with mutation (Figure 2a) or with non-synonymous mutation (Figure 2b). Also shown in Figure 2 is survival analysis of the Mariathasan cohort, in which users can select bladder cancer (BC), for APC, under the optimal expression threshold (Figure 2C) or custom grouping ratio (Figure 2D). In the Mariathasan cohort, users can also analyze the survival of BC after grouping according to Alter and Non-Alter TP53 (Figure 2E). Figure 2F shows the results of a survival analysis on WT and MT under non-synonymous mutation of TP53, in the TCGA-LUAD cohort receiving conventional treatment. The p value in the survival analysis of CAMOIP is evaluated based on a log rank test. The survival curve drawn by CAMOIP web tool also contains hazard ratio (HR) values and a 95% confidence interval (CI) for Cox analysis. In addition, CAMOIP provides the option of modifying the parameter layout, such as color, font size (including horizontal and vertical coordinate font and P value), plot width, and plot height.

**Figure 2.**
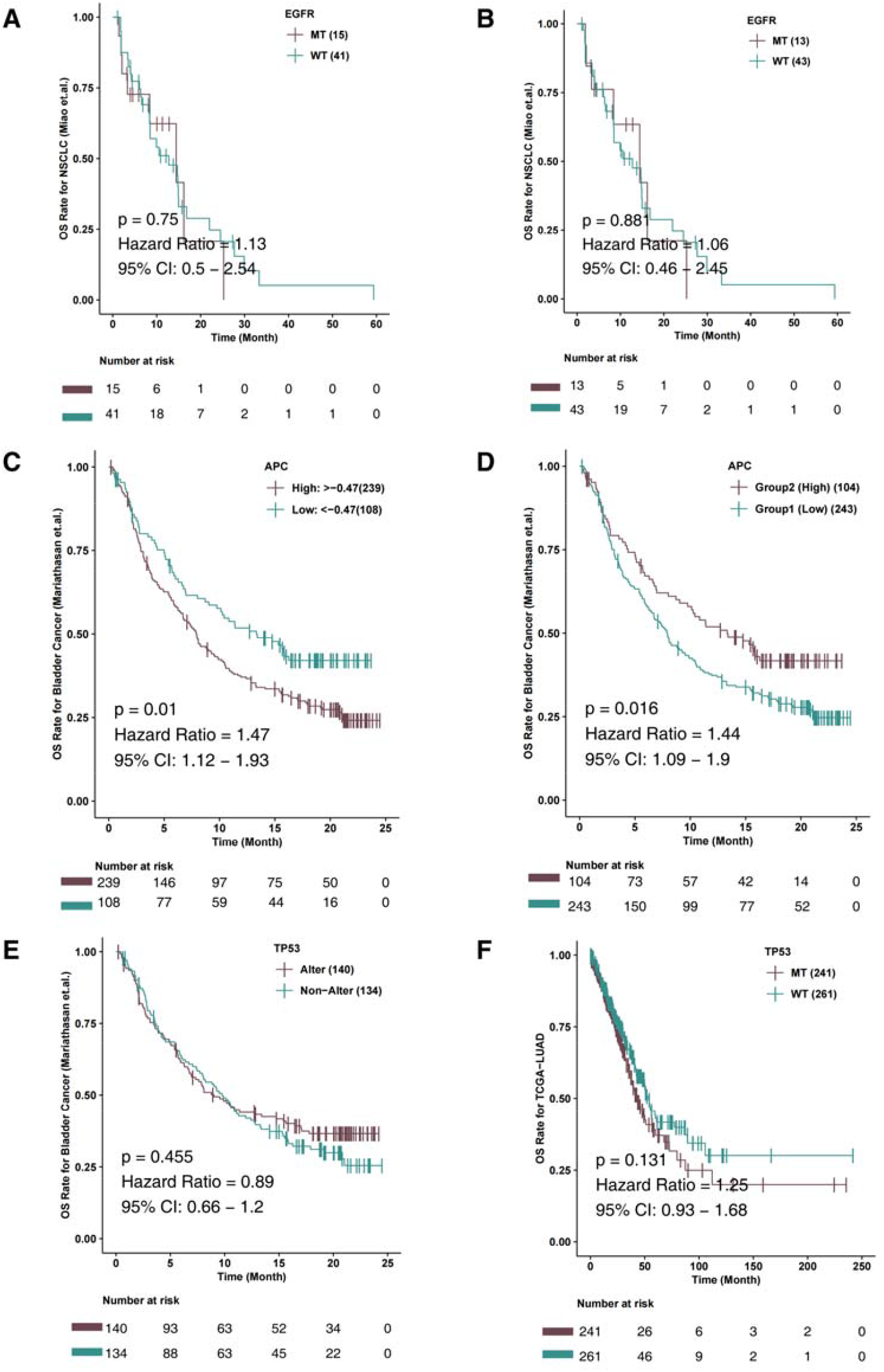
The overall survival and progression-free survival analysis of genes of interest can be displayed in the ‘Kaplan-Meier plotter’ tab. The Kaplan-Meier curve shows the comparison of OS between EGFR-MT and EGFR-WT in the Miao-NSCLC cohort using somatic mutation (A) and somatic non-synonymous mutations (B). The Kaplan-Meier curve shows the comparison of OS between APC-High and APC-Low grouped by the best cutoff (C) or certain proportion (D) in the Mariathasan-Bladder Cancer cohort using expression data. E. The Kaplan-Meier curve shows the comparison of OS between TP53-Alter and TP53-Non-Alter in the Mariathasan-Bladder Cancer cohort. F. The Kaplan-Meier curve shows the comparison of OS between TP53-MT and TP53-WT in the TCGA-LUAD cohort. HR:hazard ratio; CI: 95% confidence interval; MT: mutant; WT: wildtype; Alter: alteration; Non-Alter: non-alteration.

### GSEA

CAMOIP can perform GSEA on the expression data of different tumors according to the level of non-synonymous gene mutation or alteration, an example of which is seen in Figure 3. Users can select tumor type, specific gene and gene grouping type (mutation group: MT vs WT or gene alteration group: Alter vs Non-Alter) for subsequent analysis, and obtain an analysis result table, which can be downloaded in the form of a comma-separated values (CSV) (Figure 3A). Users can also select visualization types according to the analysis result table, such as BP plot (Figure 3B), CC plot (Figure 3C), MF plot (Figure 3D), KEGG plot (Figure 3E), REATOME plot (Figure 3f), and Heatmap (Figure 3G). The Heatmap option visualizes the expression of genes in each sample according to the specific gene set pathway. In this analysis, users can choose whether to display only the genes with a statistical difference between the two groups (display all significant genes: Yes, all significant genes or No, all genes), with red representing high gene expression, and blue representing low gene expression.

**Figure 3.**
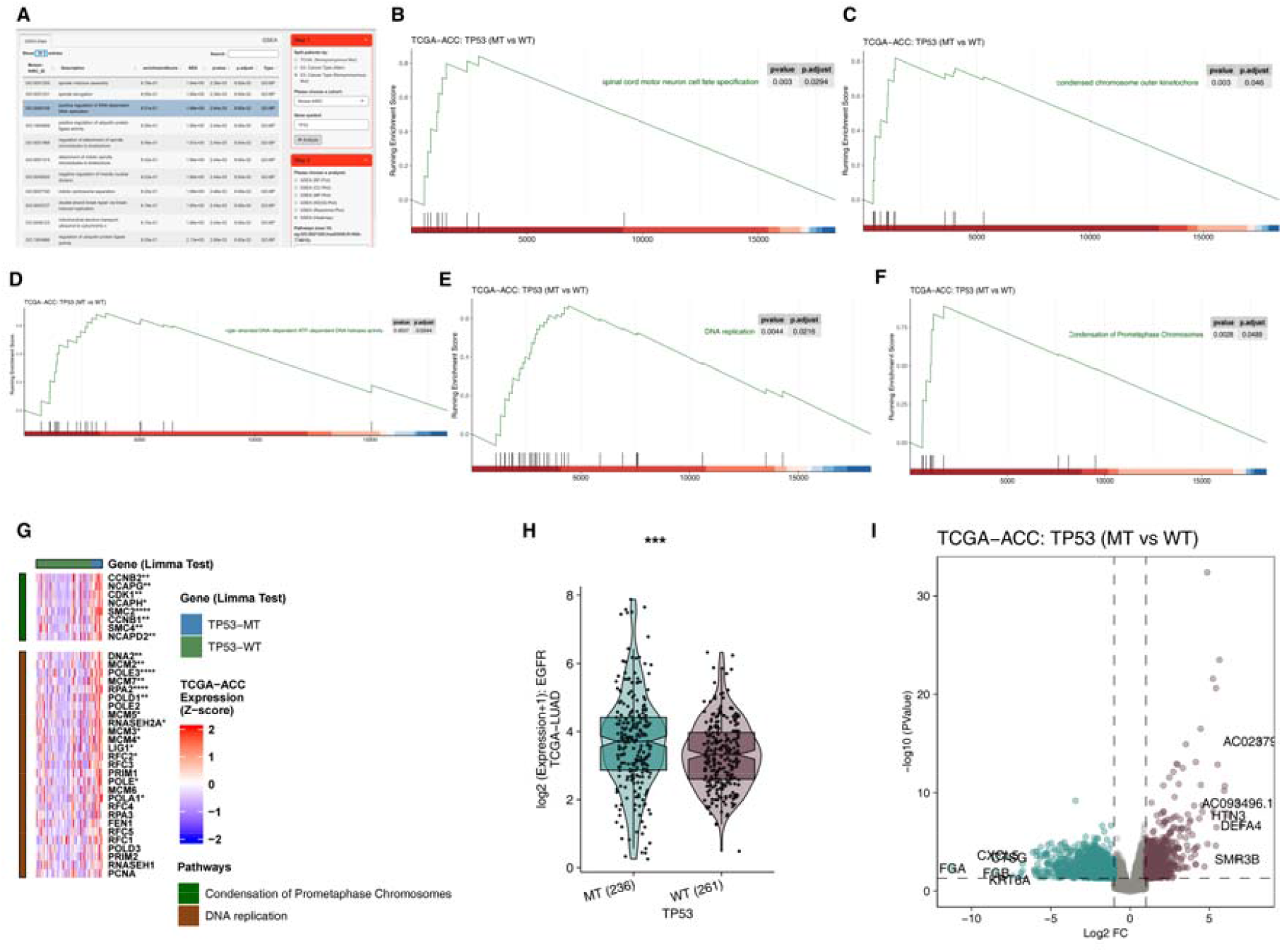
A. The result of the GSEA based on mutation type group (MT vs WT or Alter vs Non-Alter). The plot of pathways based on the BP (B), CC (C), MF (D), KEGG (E), and Reactome (F). G. Heatmap depicting the expression of genes of selected pathways in the GSEA between the MT and WT of the gene. Red indicates up-regulation, while blue indicates down-regulation. H. Comparison of gene expression between MT and WT tumors. I. The volcano plot between MT and WT tumors. GSEA: gene sets enrichment analysis; MT: mutant; WT: wildtype; Alter: alteration; Non-Alter: non-alteration; BP: biological process; CC: cellular component; MF: molecular function; KEGG: Kyoto Encyclopedia of Genes and Genomes. *: P < 0.05; **: P < 0.01; ***: P < 0.001; ****: P < 0.0001

### Expression

Under this tab, the user can select tumor type and gene expression data for differential analysis and visualization. For example, to analyze TP53 in the TCGA-LUAD cohort, the gene expression data was divided into MT and WT groups according to the non-synonymous mutation types. Under this grouping, the expression level of EGFR was tested by Mann Whitney U test and visualized by box plot (Figure 3H). In addition to the differential analysis of single gene expression, users can also perform a differential analysis on all expression data of each tumor type based on gene mutation grouping (grouped by the Bayesian LIMMA test) and visualize the results as a volcano map (Figure 3I). In addition, users can customize the output by selecting the number of top-down differential genes to label, by determining the threshold of logic and P values that defines differential genes, and by changing drawing parameters such as color, font size (including ordinate font and label font), plot width, and plot height.

### Drug Sensitivity

With this function, users can select the tumor type and gene of interest and analyze the differences in drug sensitivity. For example, users can select the GDSC-LUAD tumor cell line and compare the the half maximal inhibitory concentration (IC50) value of the drug etoposide according to TP53 mutation status (TP53-MT vs TP53-WT) (Figure 4A). According to the gene grouping (MT vs WT), the difference in sensitivity to all drugs was analyzed, the results visualized as a box plot (plot tab). This plot displays all drugs with statistical difference under grouping variables or a selected number of drugs with statistical difference (Figure 4B). The results of all drug sensitivity analyses are displayed in the All-Drug U test tab and are available to download as a CSV (Figure 4C).

**Figure 4.**
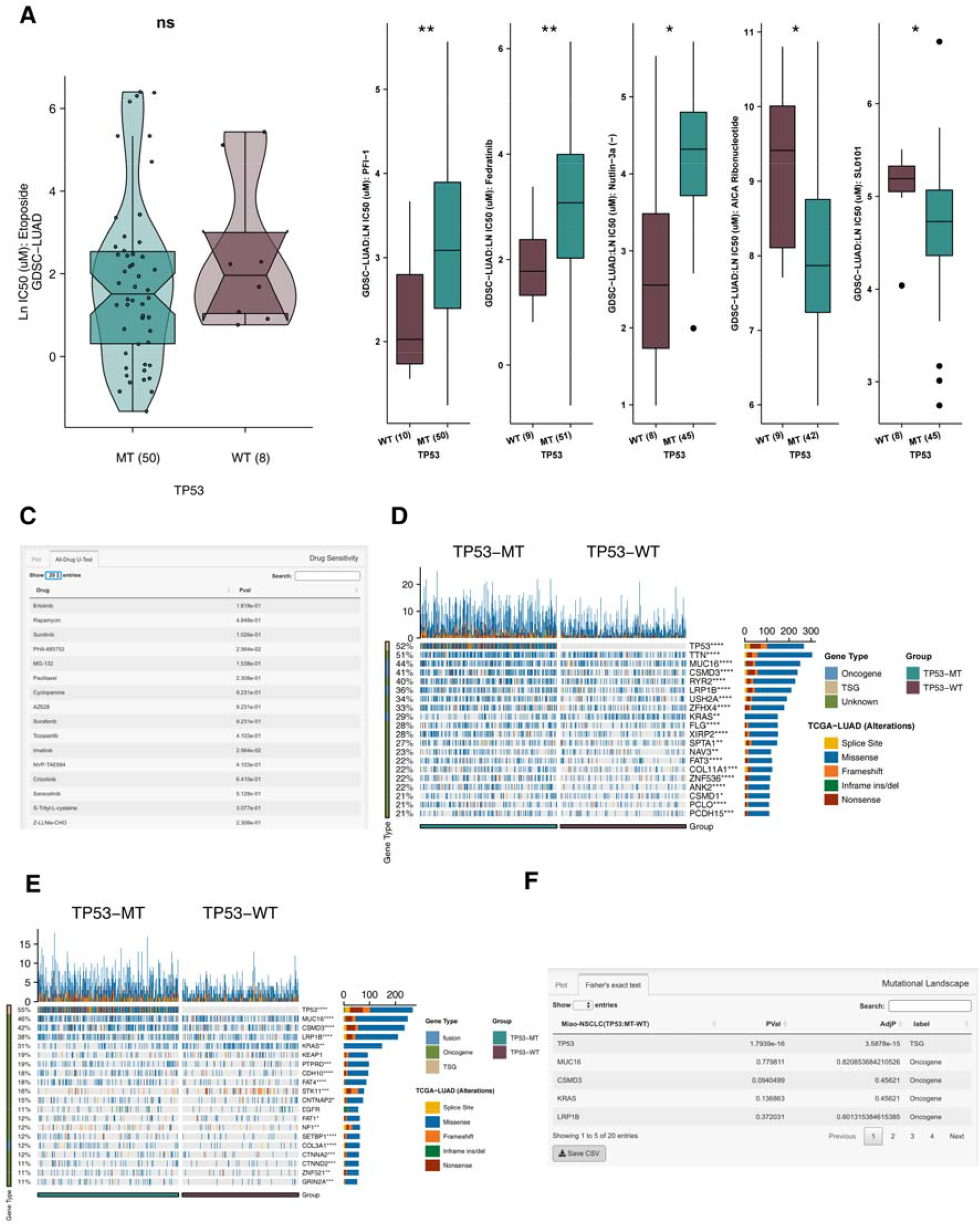
A. The comparison of the drug sensitivity (etoposide) between TP53-MT and TP53-WT in the GDSC-LUAD. B. The box plot shows the drugs with significant differences between the MT and WT groups. C. The result of drugs with significant differences between the MT and WT groups. D. The top mutated genes between TP53-MT and TP53-WT in the TCGA-LUAD cohort. E. The top mutated driver genes between TP53-MT and TP53-WT in the TCGA-LUAD cohort. F. The result of difference in mutation rate of driver genes between the MT and WT group. MT: mutant; WT: wildtype. *: P < 0.05; **: P < 0.01; ***: P < 0.001; ****: P < 0.0001

### Mutation Landscape

This function allows users to display an overview of gene mutation under different mutations or alterations groups. According to the specific tumor type and input of a specific gene, the user is able to group the non-synonymous mutation status or change the status of the gene, and further compare the difference in mutation frequency of other genes in the same cohort under the grouping variable. For example, users can select the TCGA-LUAD cohort, and according to the non-synonymous mutation status of TP53, divide the patients into TP53-MT and TP53-WT, allowing for analysis of a specific number of top genes to be compared (Figure 4D). In addition, this function provides users with a visual overview of mutations in the driver gene (Figure 4E). For example, users can select the TCGA-LUAD cohort, and for the TP53-MT and TP53-WT groups, compare the difference in mutation frequency and gene mutation type for a specific number of top driver genes (Figure 4F). Fisher’s exact test is used to obtain the p value of mutation frequency after comparison under grouping variables, and either adjusted p value (BH method) or p value can be displayed. This function also provides image modification parameters: such as color, font size (including ordinate font and label font), plot width, and plot height.

### Immune Infiltration

With this function, CAMOIP provides an opportunity for users to analyze changes over time in immune cells, immune checkpoint molecules, immune related scores, and immune related genes. An example is shown in Figure 5A, where the user has analyzed the difference in the relative proportion of immune cells for the TCGA-LUAD cohort between different TP53 non-synonymous mutation states (TP53-MT vs TP53-WT). Users can also compare the differences in expression level (Figure 5B), immune related score (Figure 5C), and immune related genes (Figure 5D), as shown for the TP53-MT and TP53-WT groups in the TCGA-LUAD cohort. Mann Whitney U test is used to calculate p values for boxplots, and the Bayes LIMMA test for heat maps. Image modification parameters such as font size, plot width, and plot height are displayed at the bottom of the right sidebar.

**Figure 5.**
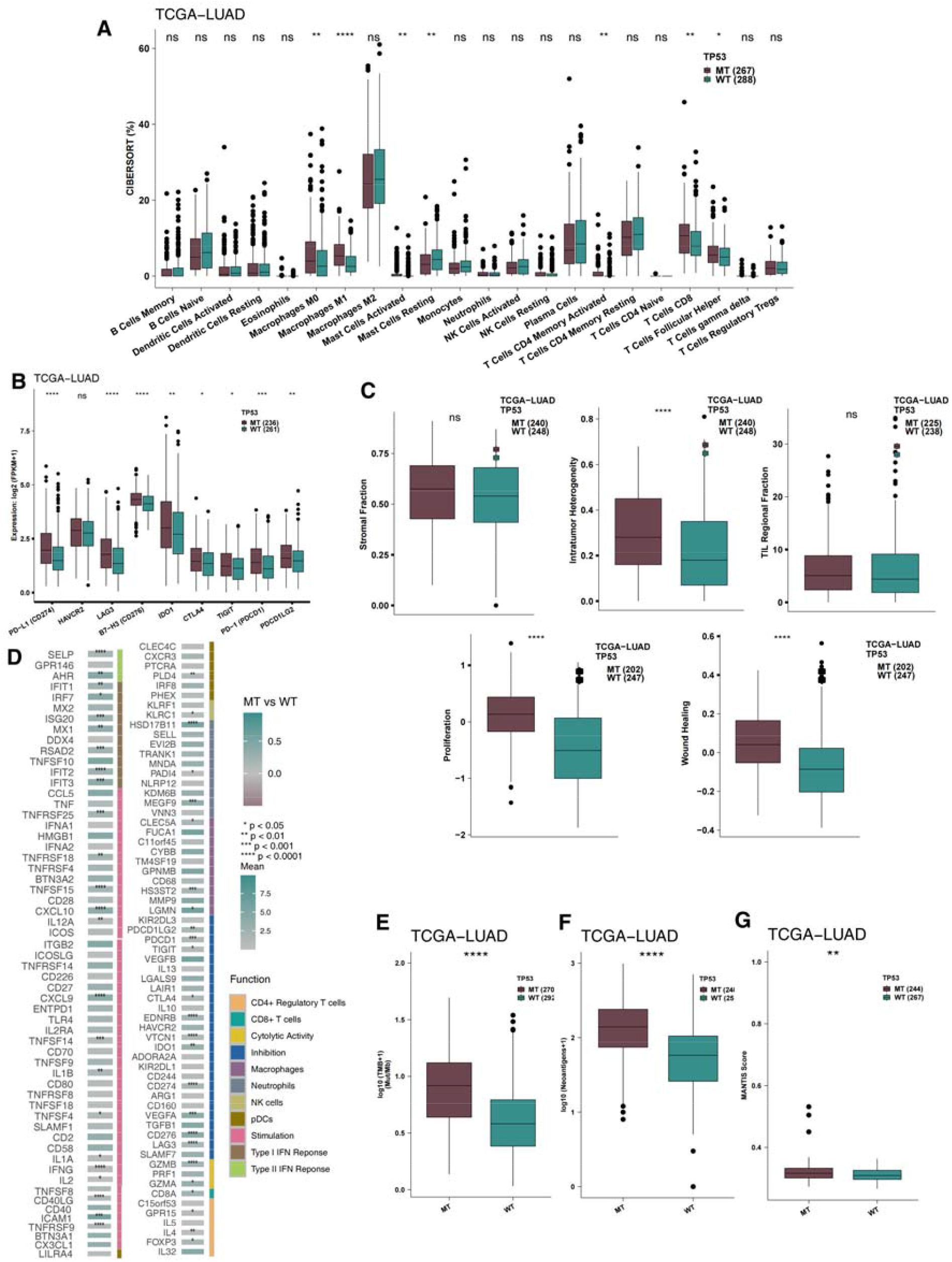
Comparison of the proportion of immune cells (A) or the expression of the immune checkpoints (B) or immune-related scores (C) or the expression of immune genes (D) between TP53-MT and TP53-WT in the TCGA-LUAD cohort. Comparison of the TMB (E) or NAL (F) or MANTIS score (G) between the MT and WT tumors. TMB: tumor mutation burdern; NAL: neoantigen loads; MT: mutant; WT: wildtype; TCGA: The Cancer Genome Atlas; LUAD: lung adenocarcinoma

### Immunogenicity

Immunogenicity is one of the most important factors influencing the response to immune checkpoint therapy, and CAMOIP provides users with a simple way to analyze it. Users can select different tumors and compare the TMB (Figure 5e), NAL (Figure 5F), and MANTIS scores (Figure 5g) of each. Mann Whitney U test is used for the analysis, and the results are visualized as a box diagram.

### Availability of results

After the user submits their request, CAMOIP provides the user with vector images, bitmaps and tables. The results provided by CAMOIP can be directly published by clicking the PNG and PDF buttons below images to download the image file in the form of a PNG or PDF, and clicking CSV at the bottom of tables to download the file in the form of a CSV. PDF image files can be further adjusted in Adobe Illustrator. Examples are provided for each tab in the example page included with CAMOIP.

### Documentation

The “Help” submenu under the “Doc” page provides the exact definition and explanation of all parameters in each tab. The “Q & A” submenu under the “Doc” page lists common problems and solutions that users may come across while using the CAMOIP web tool.

## DISCUSSION

As an interactive web tool, CAMOIP mainly uses information from clinical and TCGA cohorts on expression, mutation, and clinical prognosis data (for both immunotherapy and conventional treatment). CAMOIP provides for biologists, without the need for programming technology or expertise, the ability to screen for immune checkpoint therapy biomarkers as well as the mechanism of the biomarkers associated with changes in the tumor immune microenvironment. In doing so, the CAMOIP tool allows users to easily evaluate a gene mutation or change, such as whether a certain gene expression level is related to the prognosis of tumor patients after receiving immune checkpoint treatment, and to verify the correlation mechanism between immunotherapy biomarkers and treatment response. For example, TP53 can be used as a biomarker for the prognosis of immunotherapy cohort Samstein-BLCA, which can be easily investigated by users in CAMOIP(36). In addition, users can find that ZFHX3 is associated with significantly longer OS time in the immunotherapy cohort of Samstein-NSCLC(22). The CARD11 gene can also be quickly identified as a gene significantly associated with the prognosis of skin cutaneous melanoma (SKCM) patients receiving immunotherapy(37).

In addition to screening biomarkers for the prognosis of immunotherapy patients with different tumors, users can also use the GSEA, expression, drug sensitivity, mutation landscape, immune infection, and immunology tabs to explore the specific mechanism of the correlation between the selected biomarkers and immunotherapy. At the same time, the custom drawing parameters of CAMOIP allow users great freedom to customize the visual output, for example, by changing the color, changing the font size, changing the form of display for p values, modifying the plot height, and adjusting the image width.

CAMOIP is a time-saving and intuitive web-based tool that provides users, especially biologists with no background knowledge of computer programming, with a powerful tool for mining biomarkers and exploring the follow-up mechanism of immunotherapy prognosis. With continuous user feedback and further enhancement, CAMOIP has the potential to become an integral part of routine data analysis for experimental biologists.

## DATA AVAILABILITY

R studio is a language and environment for statistical computing available in https://www.r-project.org/.

Shiny is an open source collaborative initiative available in the GitHub repository (https://github.com/rstudio/shiny).

TCGAbiolinks is an open source collaborative initiative available in the GitHub repository (https://github.com/BioinformaticsFMRP/TCGAbiolinks).

## SUPPLEMENTARY DATA

None

## ACKNOWLEDGEMENT

Special thanks to the English language polishing contributions from TopScience Editing.

## FUNDING

None

## CONFLICT OF INTEREST

The authors declare that the research was conducted in the absence of any commercial or financial relationships that could be construed as a potential conflict of interest.

## Notes

### Competing Interest Statement

The authors have declared no competing interest.

## REFERENCES

1. O’Donnell, J.S., Teng, M.W.L. and Smyth, M.J. (2019) Cancer immunoediting and resistance to T cell-based immunotherapy. Nat. Rev. Clin. Oncol., 16, 151–167.

2. Huang, X., Tang, T., Zhang, G., Hong, Z., Xu, J., Yadav, D.K., Bai, X. and Liang, T. (2020) Genomic investigation of co-targeting tumor immune microenvironment and immune checkpoints in pan-cancer immunotherapy. NPJ Precis. Oncol., 4, 29.

3. Niu, Y., Lin, A., Luo, P., Zhu, W., Wei, T., Tang, R., Guo, L. and Zhang, J. (2020) Prognosis of Lung Adenocarcinoma Patients With NTRK3 Mutations to Immune Checkpoint Inhibitors. Front. Pharmacol., 11, 1213.

4. Lin, A., Zhang, H., Hu, X., Chen, X., Wu, G., Luo, P. and Zhang, J. (2020) Age, sex, and specific gene mutations affect the effects of immune checkpoint inhibitors in colorectal cancer. Pharmacol. Res., 159, 105028.

5. Samstein, R.M., Lee, C.-H., Shoushtari, A.N., Hellmann, M.D., Shen, R., Janjigian, Y.Y., Barron, D.A., Zehir, A., Jordan, E.J., Omuro, A., et al. (2019) Tumor mutational load predicts survival after immunotherapy across multiple cancer types. Nat. Genet., 51, 202–206.

6. Prat, A., Navarro, A., Paré, L., Reguart, N., Galván, P., Pascual, T., Martínez, A., Nuciforo, P., Comerma, L., Alos, L., et al. (2017) Immune-Related Gene Expression Profiling After PD-1 Blockade in Non-Small Cell Lung Carcinoma, Head and Neck Squamous Cell Carcinoma, and Melanoma. Cancer Res., 77, 3540–3550.

7. Rizvi, H., Sanchez-Vega, F., La, K., Chatila, W., Jonsson, P., Halpenny, D., Plodkowski, A., Long, N., Sauter, J.L., Rekhtman, N., et al. (2018) Molecular Determinants of Response to Anti-Programmed Cell Death (PD)-1 and Anti-Programmed Death-Ligand 1 (PD-L1) Blockade in Patients With Non-Small-Cell Lung Cancer Profiled With Targeted Next-Generation Sequencing. J. Clin. Oncol. Off. J. Am. Soc. Clin. Oncol., 36, 633–641.

8. Hwang, S., Kwon, A.-Y., Jeong, J.-Y., Kim, S., Kang, H., Park, J., Kim, J.-H., Han, O.J., Lim, S.M. and An, H.J. (2020) Immune gene signatures for predicting durable clinical benefit of anti-PD-1 immunotherapy in patients with non-small cell lung cancer. Sci. Rep., 10, 643.

9. Van Allen, E.M., Miao, D., Schilling, B., Shukla, S.A., Blank, C., Zimmer, L., Sucker, A., Hillen, U., Foppen, M.H.G., Goldinger, S.M., et al. (2015) Genomic correlates of response to CTLA-4 blockade in metastatic melanoma. Science, 350, 207–211.

10. Ulloa-Montoya, F., Louahed, J., Dizier, B., Gruselle, O., Spiessens, B., Lehmann, F.F., Suciu, S., Kruit, W.H.J., Eggermont, A.M.M., Vansteenkiste, J., et al. (2013) Predictive gene signature in MAGE-A3 antigen-specific cancer immunotherapy. J. Clin. Oncol. Off. J. Am. Soc. Clin. Oncol., 31, 2388–2395.

11. Hugo, W., Zaretsky, J.M., Sun, L., Song, C., Moreno, B.H., Hu-Lieskovan, S., Berent-Maoz, B., Pang, J., Chmielowski, B., Cherry, G., et al. (2016) Genomic and Transcriptomic Features of Response to Anti-PD-1 Therapy in Metastatic Melanoma. Cell, 165, 35–44.

12. Auslander, N., Zhang, G., Lee, J.S., Frederick, D.T., Miao, B., Moll, T., Tian, T., Wei, Z., Madan, S., Sullivan, R.J., et al. (2018) Robust prediction of response to immune checkpoint blockade therapy in metastatic melanoma. Nat. Med., 24, 1545–1549.

13. Riaz, N., Havel, J.J., Makarov, V., Desrichard, A., Urba, W.J., Sims, J.S., Hodi, F.S., Martín-Algarra, S., Mandal, R., Sharfman, W.H., et al. (2017) Tumor and Microenvironment Evolution during Immunotherapy with Nivolumab. Cell, 171, 934–949.e16.

14. Sui, S., An, X., Xu, C., Li, Z., Hua, Y., Huang, G., Sui, S., Long, Q., Sui, Y., Xiong, Y., et al. (2020) An immune cell infiltration-based immune score model predicts prognosis and chemotherapy effects in breast cancer. Theranostics, 10, 11938–11949.

15. Dolled-Filhart, M., Locke, D., Murphy, T., Lynch, F., Yearley, J.H., Frisman, D., Pierce, R., Weiner, R., Wu, D. and Emancipator, K. (2016) Development of a Prototype Immunohistochemistry Assay to Measure Programmed Death Ligand-1 Expression in Tumor Tissue. Arch. Pathol. Lab. Med., 140, 1259–1266.

16. Mino-Kenudson, M. (2016) Programmed cell death ligand-1 (PD-L1) expression by immunohistochemistry: could it be predictive and/or prognostic in non-small cell lung cancer? Cancer Biol. Med., 13, 157–170.

17. Ghatalia, P. and Plimack, E. (2019) Biomarkers for neoadjuvant checkpoint blockade response in urothelial cancer. Nat. Med., 25, 1650–1651.

18. Yarchoan, M., Hopkins, A. and Jaffee, E.M. (2017) Tumor Mutational Burden and Response Rate to PD-1 Inhibition. N. Engl. J. Med., 377, 2500–2501.

19. Lin, A., Qiu, Z., Zhang, J. and Luo, P. (2021) Effect of NCOR1 Mutations on Immune Microenvironment and Efficacy of Immune Checkpoint Inhibitors in Patient with Bladder Cancer. Front. Immunol., 12, 630773.

20. Endris, V., Buchhalter, I., Allgäuer, M., Rempel, E., Lier, A., Volckmar, A.-L., Kirchner, M., von Winterfeld, M., Leichsenring, J., Neumann, O., et al. (2019) Measurement of tumor mutational burden (TMB) in routine molecular diagnostics: in silico and real-life analysis of three larger gene panels. Int. J. cancer, 144, 2303–2312.

21. Meléndez, B., Van Campenhout, C., Rorive, S., Remmelink, M., Salmon, I. and D’Haene, N. (2018) Methods of measurement for tumor mutational burden in tumor tissue. Transl. lung cancer Res., 7, 661–667.

22. Zhang, J., Zhou, N., Lin, A., Luo, P., Chen, X., Deng, H., Kang, S., Guo, L., Zhu, W. and Zhang, J. (2021) ZFHX3 mutation as a protective biomarker for immune checkpoint blockade in non-small cell lung cancer. Cancer Immunol. Immunother., 70, 137–151.

23. Warth, A., Körner, S., Penzel, R., Muley, T., Dienemann, H., Schirmacher, P., von Knebel-Doeberitz, M., Weichert, W. and Kloor, M. (2016) Microsatellite instability in pulmonary adenocarcinomas: a comprehensive study of 480 cases. Virchows Arch., 468, 313–319.

24. Lin, A., Zhang, J. and Luo, P. (2020) Crosstalk Between the MSI Status and Tumor Microenvironment in Colorectal Cancer. Front. Immunol., 11, 2039.

25. Goldman, M., Craft, B., Brooks, A., Zhu, J. and Haussler, D. (2018) The UCSC Xena Platform for cancer genomics data visualization and interpretation. biorxiv.

26. Gao, J., Aksoy, B.A., Dogrusoz, U., Dresdner, G., Gross, B., Sumer, S.O., Sun, Y., Jacobsen, A., Sinha, R., Larsson, E., et al. (2013) Integrative analysis of complex cancer genomics and clinical profiles using the cBioPortal. Sci. Signal., 6, pl1.

27. Uhlén, M., Fagerberg, L., Hallström, B.M., Lindskog, C., Oksvold, P., Mardinoglu, A., Sivertsson, Å., Kampf, C., Sjöstedt, E., Asplund, A., et al. (2015) Proteomics. Tissue-based map of the human proteome. Science, 347, 1260419.

28. Petryszak, R., Keays, M., Tang, Y.A., Fonseca, N.A., Barrera, E., Burdett, T., Füllgrabe, A., Fuentes, A.M.-P., Jupp, S., Koskinen, S., et al. (2016) Expression Atlas update--an integrated database of gene and protein expression in humans, animals and plants. Nucleic Acids Res., 44, D746–52.

29. Tang, Z., Li, C., Kang, B., Gao, G., Li, C. and Zhang, Z. (2017) GEPIA: a web server for cancer and normal gene expression profiling and interactive analyses. Nucleic Acids Res., 45, W98–W102.

30. Tang, Z., Kang, B., Li, C., Chen, T. and Zhang, Z. (2019) GEPIA2: an enhanced web server for large-scale expression profiling and interactive analysis. Nucleic Acids Res., 47, W556–W560.

31. Chang, W., Cheng, J., Allaire, J.J., Xie, Y. and McPherson, J. (2015) Package ‘shiny’. See http://citeseerx.ist.psu.edu/viewdoc/download.

32. Edgar, R., Domrachev, M. and Lash, A.E. (2002) Gene Expression Omnibus: NCBI gene expression and hybridization array data repository. Nucleic Acids Res., 30, 207–210.

33. Colaprico, A., Silva, T.C., Olsen, C., Garofano, L., Cava, C., Garolini, D., Sabedot, T.S., Malta, T.M., Pagnotta, S.M., Castiglioni, I., et al. (2016) TCGAbiolinks: an R/Bioconductor package for integrative analysis of TCGA data. Nucleic Acids Res., 44, e71.

34. Jensen, M.A., Ferretti, V., Grossman, R.L. and Staudt, L.M. (2017) The NCI Genomic Data Commons as an engine for precision medicine. Blood, 130, 453–459.

35. Yang, W., Soares, J., Greninger, P., Edelman, E.J., Lightfoot, H., Forbes, S., Bindal, N., Beare, D., Smith, J.A., Thompson, I.R., et al. (2013) Genomics of Drug Sensitivity in Cancer (GDSC): a resource for therapeutic biomarker discovery in cancer cells. Nucleic Acids Res., 41, D955–61.

36. Lyu, Q., Lin, A., Cao, M., Xu, A., Luo, P. and Zhang, J. (2020) Alterations in TP53 Are a Potential Biomarker of Bladder Cancer Patients Who Benefit From Immune Checkpoint Inhibition. Cancer Control, 27, 1073274820976665.

37. Wen, Y., Lin, A., Zhu, W., Wei, T., Luo, P., Guo, L. and Zhang, J. (2021) Catenin Alpha-2 Mutation Changes the Immune Microenvironment in Lung Adenocarcinoma Patients Receiving Immune Checkpoint Inhibitors. Front. Pharmacol., 12, 645862.

